# Climate risk to European fisheries and coastal communities

**DOI:** 10.1101/2020.08.03.234401

**Authors:** Mark R. Payne, Manja Kudahl, Georg H. Engelhard, Myron A. Peck, John K. Pinnegar

**Author notes:** Corresponding author. Mark R. Payne, National Institute of Aquatic Resources (DTU-Aqua), Technical University of Denmark, 2800 Kgs. Lyngby Denmark.

## Abstract

With the majority of the global human population living in coastal regions, correctly characterising the climate risk that ocean-dependent communities and businesses are exposed to is key to prioritising the finite resources available to support adaptation. We apply a climate risk analysis across the European fisheries sector for the first time to identify the most at-risk fishing fleets and coastal regions and then link the two analyses together. We employ a novel approach combining biological traits with physiological metrics to differentiate climate hazards between 556 populations of fish and use these to assess the relative climate risk for 380 fishing fleets and 105 coastal regions in Europe. Countries in southeast Europe as well as the UK have the highest risks to both fishing fleets and coastal regions overall while, in other countries, the risk-profile is greater at either the fleet level or at the regional level. European fisheries face a diversity of challenges posed by climate change and climate adaptation, therefore, needs to be tailored to each country, region and fleet’s specific situation. Our analysis supports this process by highlighting where and what adaptation measures might be needed and informing where policy and business responses could have the greatest impact.

**Significance Statement:** We present a novel climate risk analysis for i) 105 ocean-dependent communities and ii) 380 fishing fleets in Europe. Our unique approach provides a perspective over the climate risks in this diverse and populous continent that is unprecedented in both its breadth and detail. We show that countries in southeast Europe as well as the UK have the highest climate risk overall, both in terms of fishing fleets and coastal communities. Substantial variation in climate risk is seen even within countries, emphasizing that climate adaptation interventions need to be tailored to the specific characteristics of the fleet or community. A focus on sustainable fisheries management and diversification of fish portfolios can reduce climate risks across the board.

## Main Text

The ocean provides human societies with a wide variety of goods and services, ranging from food and employment to climate regulation and cultural nourishment (Hassan *et al*., 2005). Climate change is already shifting the abundance, distribution, productivity and phenology of living marine resources (Poloczanska *et al*., 2013; FAO, 2018; Phillips and Pérez-Ramírez, 2018), thereby impacting many of the ecosystem services upon which society depends (IPCC, 2019). These impacts, however, are not being experienced uniformly by human society but depend on the characteristics and context of the community or business affected. Raising awareness and understanding the risk to human systems is therefore a critical first step (Allison *et al*., 2009) to developing and prioritising appropriate adaptation options in response to the challenges of the climate crisis (Johnson *et al*., 2016).

Over the past decades, climate risk assessments (CRAs) and climate vulnerability assessments (CVAs) have been developed to identify and prioritise adaptation needs. The approach, developed by the Intergovernmental Panel on Climate Change (IPCC), has shifted over time from a focus on “vulnerability” to a focus on “risk” (Oppenheimer *et al*., 2014), in part due to criticisms of the negative framing that “vulnerability” implies (Connelly *et al*., 2018). The modern CRA framework (Cardona *et al*., 2012) considers risk as the intersection of hazard, exposure and vulnerability (Table 1). CVAs, and more recently CRAs, have been applied widely in the marine realm, for example in coastal communities in northern Vietnam (Adger, 1999), Kenya (Cinner *et al*., 2013) and the USA (Colburn *et al*., 2016), at the national level across coastal areas of the USA (Ekstrom *et al*., 2015; Hare *et al*., 2016) and Australia (Pecl *et al*., 2014; Fulton *et al*., 2017), across regions such as Pacific island nations (Bell *et al*., 2011; Barsley *et al*., 2013) and globally (Allison *et al*., 2009; Barange *et al*., 2014; Blasiak *et al*., 2017). Several ‘best practice’ guides have also been developed (Brugere and De Young, 2015; Johnson *et al*., 2016).

**Table 1.** Definitions of terms, as used in the context of this Climate Risk Analysis. These definitions are adapted for the present study from those used in the most recent IPCC report (IPCC, 2019).

| Term | Definition used here |
| --- | --- |
| **Climate risk** | The potential for climate change to have adverse consequences for human systems, specifically for European coastal regions and fishing fleets. |
| **Hazard** | The potential for, and severity of, climate change impacts on the unit of interest (i.e. fish and shellfish populations). Here we focus explicitly on negative impacts, following from the definition of risk as being an adverse consequence. |
| **Exposure** | The sensitivity of a region or fishing fleet to the climate hazard, i.e. the likelihood of being affected by changes in the living marine resources. |
| **Vulnerability** | The ability of a region or fleet to anticipate or respond to changes induced by climate hazards and to minimise, cope with and recover from the consequences. High adaptive capacity gives low vulnerability. |

CRAs and CVAs covering European waters are, however, notable by their absence from this list. This is surprising given that European waters provide over one-eighth of the world’s total marine fisheries catches (FAO, 2020), and have witnessed many well-documented changes in fish abundance and distribution in response to climate change (Perry *et al*., 2005; Peck and Pinnegar, 2018; Baudron *et al*., 2020). The lack of attention to climate risk in European fisheries may be due, in part, to the previous results of global CVAs (Allison *et al*., 2009) that ranked European countries as having low vulnerabilities (their relative affluence giving high ‘adaptive capacity’ in these analyses). Yet the European region poses unique challenges when assessing climate risks due to the wide range of species, biogeographical zones and habitats linked by intertwined management structures. Fishing techniques and the scale of fisheries also vary widely, from large fleets of small vessels in the Mediterranean Sea (STECF, 2018) to some of the largest fishing vessels in the world (e.g. the 144-m long *Annelies Ilena*). Furthermore, although fisheries contribute very little to national GDP, food or income-security for most European countries (Peck and Pinnegar, 2018), in specific communities and regions fishing is the mainstay of employment (Natale *et al*., 2013). Adapting European fisheries to a changing climate, therefore, requires the development of robust analyses capable of assessing the climate risk across this extremely diverse continent.

We conducted a CRA across the European marine fisheries sector that is globally unprecedented in its span and detail, estimating the climate risk of i) coastal regions and ii) fishing fleets in linked analyses. Our analyses spanned more than 50 degrees of latitude from the Black Sea to the Arctic and encompass the United Kingdom, Norway, Iceland and Turkey in addition to the 22 coastal nations of the European Union. We developed an entirely novel approach that distinguishes fine-scale geographical differences in the climate hazard of fish and shellfish populations and, hence, the climate risk to both European coastal regions and fishing fleets. Uniquely, since both CRAs were based on the same underlying climate hazard, these analyses could be combined to compare the relative importance of this hazard to fleets and coastal regions within a country.

### Coastal-Region Climate Risk Analysis

Our novel index of climate hazard was derived from the biological traits of the species being harvested, together with modelled distribution data. Species trait data were gathered for 157 fish and shellfish species harvested in European waters, representing 90.3% of the total value of landings in Europe and at least 78% (and typically more than 90%) of national value. Uniquely, we accounted for the expected large variation in climate hazard throughout a species range (i.e. from the cold to warm edges of the distribution) by focusing on “populations” (i.e. a single species in a single FAO sub-area). Population-level climate hazards were then defined based on estimates of the thermal-safety margin (TSM) (*sensu* Deutsch *et al*., 2008; Sunday *et al*., 2014; Dahlke *et al*., 2020) between the temperature in that subregion and the empirically-derived upper thermal preference of the species (Kesner-Reyes *et al*., 2012). Climate hazards were calculated for 556 “populations” in 23 FAO subareas, based on the TSM of the population and the inherent traits of the species (Cheung *et al*., 2005; Hare *et al*., 2016; Jones and Cheung, 2018).

We then calculated the climate risk for 105 coastal regions across 26 countries in the European continent (Figure 1). Population-level climate hazards of fish were integrated to regions, weighted by the relative value of landings in that region. We defined exposure metrics based on the diversity and dominance (Cline *et al*., 2017; Pinnegar *et al*., 2019) of these landings, and vulnerability based on regional socio-economic metrics (Allison *et al*., 2009). We focused our analysis on coastal regions, as these are the communities most directly dependent on the ocean: regions far from the sea but within a coastal nation were explicitly excluded (e.g. Bavaria in the south of Germany).

**Figure 1.**
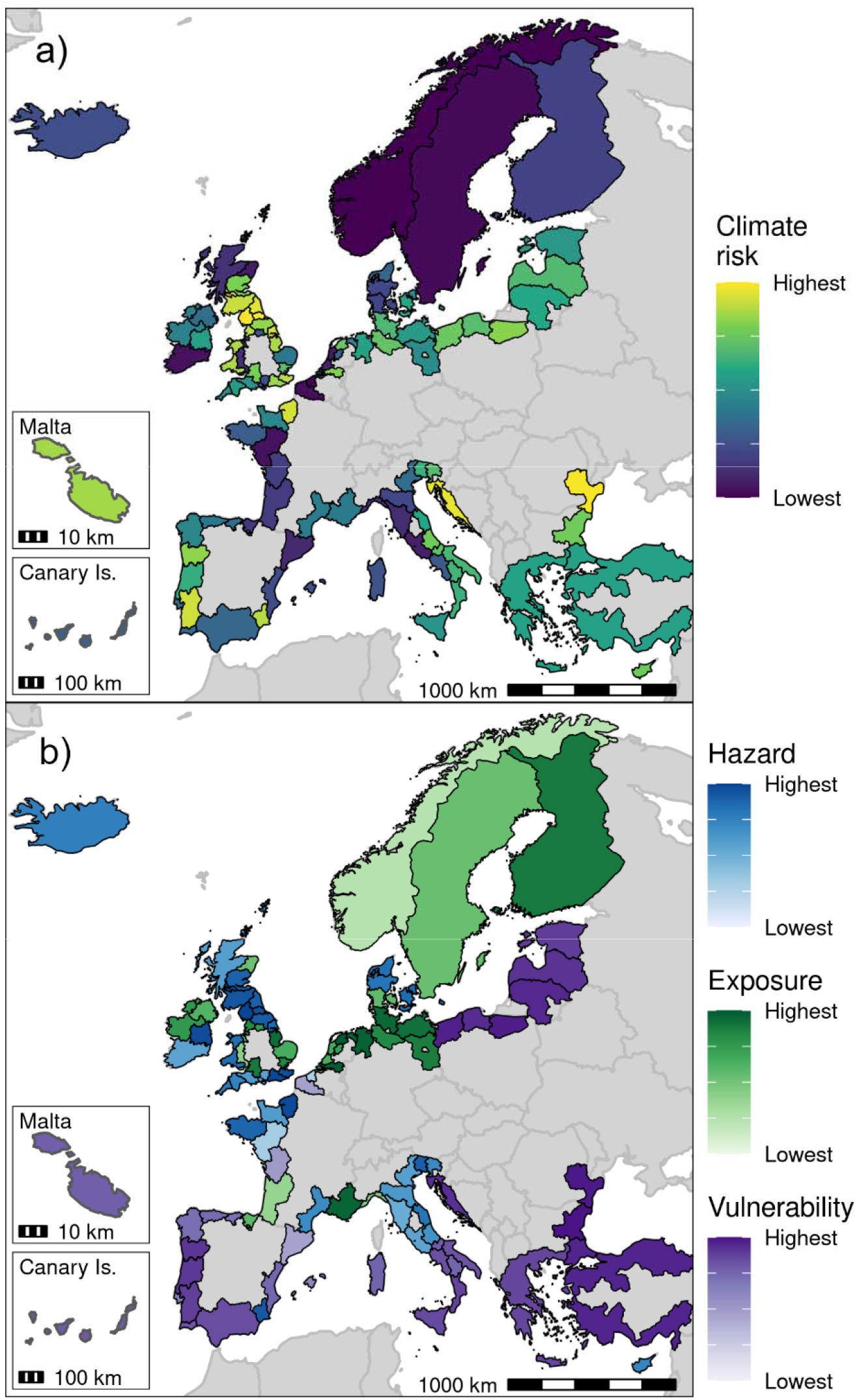
Climate risk of European regions. Maps show a) the combined climate risk ranking for each region and b) the individual component (blue: hazard, green: exposure, purple: vulnerability) making the largest contribution to the combined risk. Colour scales on both panels are linear in the ranking of the corresponding score, but are presented without values, as they have little direct meaning. National borders are also shown for reference. Insets at bottom-left of each panel show small regions. Maps showing the hazard, exposure and vulnerability for each coastal region are included in the supplementary material (Figure S1).

**Figure 2.**
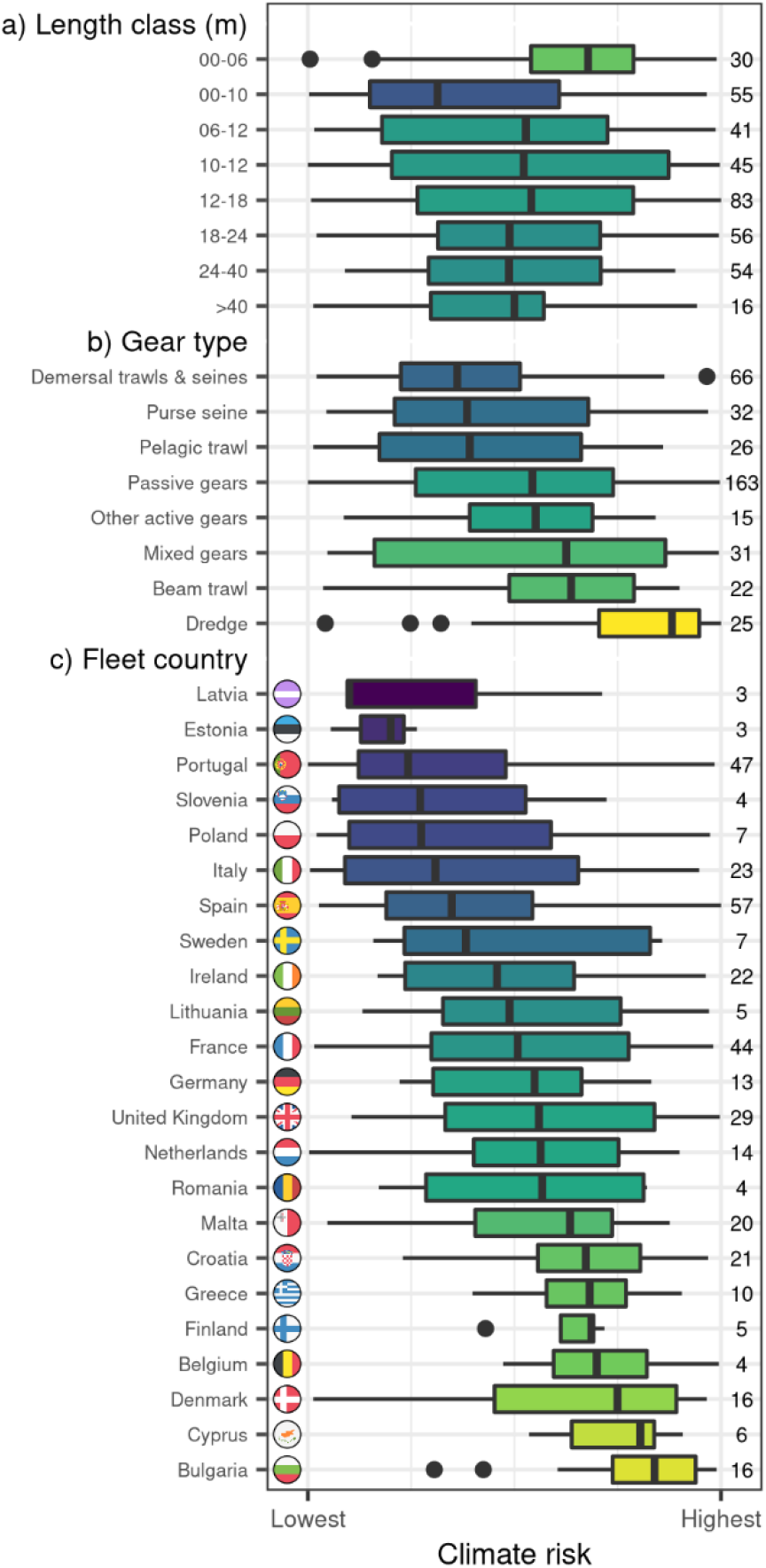
Climate risk of European fleet segments. The climate risk ranking across 380 fleet segments is plotted as a function of a) the size range of the vessels (m), b) the gear type employed (sorted by median risk) and c) the country of origin of the fleet (sorted by median risk). Risk ranking is represented on a linear scale from highest to lowest: the absolute values of risk are not shown, as they have little direct meaning. The distribution of risk is shown as a boxplot, where the vertical line is the median, the box corresponds to the interquartile range (IQR), and the whiskers cover all points less than 1.5 times the IQR from the box. Outliers are plotted as points. Boxes are coloured based on the median climate risk for that category. The number of fleet segments in each class is shown at right. Note that EU definitions of small length classes (less than 12m) vary between individual countries and therefore have a degree of overlap. Specific gear codes are aggregated here to broader-scale categories of “Gear Types” to ensure comparability between Atlantic and Mediterranean fisheries (Table S4).

The analysis revealed appreciable variation in the climate risk within the European continent and even within a single country (Figure 1a). In the United Kingdom, for example, climate risk was greatest in the north of England, while Scotland and the south of England had the least risk. Indeed, six of the 10 regions with the highest climate risk, including the overall top region (Tees Valley & Durham), were in the UK (Table S8). These results were strongly influenced by high hazard scores for the species landed in these regions (Figure 1b), combined with high vulnerability due to low GDP per capita in some of these regions.

Larger-scale patterns in climate risk were also apparent. South-east Europe stood out with consistently high climate risk, with coastal Romania and Croatia in the top five. Both countries had high vulnerability scores due to low GDP per capita of their coastal regions (Figure 1b, Figure S1), and high exposure scores due to fisheries that target only a few species (e.g. the value of Romania’s fisheries is more than 70% veined rapa whelk, *Rapana venosa*). Many northern European nations, including Belgium, the Netherlands and Scandinavian nations had relatively low climate risks due to their wealth (high GDP per capita), diverse fisheries and the relatively low climate hazard of the fish populations targeted (Figure S1).

These overall climate-risk scores were heavily influenced by the elements (hazard, exposure or vulnerability) that comprise their risk profile (Figure 1b; Figure S1). The climate risk profiles of south-east Europe, the Iberian peninsula and some regions on the south coast of the Baltic Sea were dominated by the vulnerability dimension, reflecting the low GDP per capita of these regions. For the most part, exposure scores were important in Northern Europe and in Scandinavia, reflecting the narrower range of species landed compared to the Mediterranean region. The climate risk of Iceland, the UK, and parts of France or northern Italy, on the other hand, were dominated by the climate hazard component, i.e. the traits and thermal preferences of the species targeted. The relative contributions of the individual components are critical to understanding the climate risk of each country and the suitability of particular adaptation responses.

### Fleet Segment Climate Risk Analysis

The risks associated with climate change will also be felt directly by fishing vessels and fleets in addition to regions: we therefore performed a second CRA to examine the climate risk of European fishing fleets. As the basis for this analysis, we followed the EU definition of a “fleet segment” based on the size classes of the vessels, the country of registration, the gear being used and the geographical region being fished (Atlantic or Mediterranean) (STECF, 2018). We integrated climate hazards at the fish population level up to the fleet segment level based on the composition of landings by value of that fleet, while we based exposure on the diversity and dominance of landings and vulnerability on the net profitability of the fleet. Coverage of our analysis at this fleet segment level was less than at the national level: nevertheless, we still covered 75% or more of total fishery catch value for more than 70% of the 380 fleet segments within the EU and UK.

The smallest class of vessels (0-6m) had an appreciably higher climate risk than all other size classes (Figure 2a). For the most part, these fleets operated in the Mediterranean region, particularly in Croatia, Bulgaria, France, Malta and Greece (Table S9). This result reflected, in part, the higher climate risk of stocks in this area, but was also driven by the poor profitability (and therefore higher vulnerability) of these fleets. On the other hand, the high catch diversity of these fleets reduced exposure and helped reduce their net climate risk.

Systematic differences in climate risk were also seen among gear types (Figure 2b), where dredgers had the highest climate risk. These fleets generally targeted populations with high climate hazards and had low species diversity in their catches (giving high exposure): good profitability, on the other hand, lowered their vulnerability and somewhat reduced overall risk (Table S9). Fleets using pelagic and demersal trawls together with purse seine fleets had the lowest climate risks, primarily due to the low hazard associated with the species on which they fish.

The strongest differentiation in climate risk between fleet segments was at the national level (Figure 2c). A clear cluster of high climate risk fleet segments could be seen in south-east Europe, particularly in Croatia, Greece, Bulgaria, Cyprus and Romania (Figure S2). The risk profiles underlying each of these cases, however, were quite different, emphasising the need to understand the components in detail. Greek and Cypriot fleets had high climate risks due to poor profitability and, therefore, high vulnerability, while Bulgarian and Romanian fleets active in the Black Sea had extremely low catch diversities, giving them unusually high exposures (Table S9). It is also important to note that there was substantial variation among fleets within a country. For example, two of the five most at-risk fleets (including the most at risk) were Spanish (Table S9), even though the national level median for Spain was amongst the lowest in Europe. A detailed examination of the individual elements of the risk-profile is therefore critical to understanding the underlying factors responsible for these results.

### Comparative Analysis

A strength of the analysis performed here is that the results of the region and fleet CRAs could be directly compared. While the regions and fleets were exposed to the same base set of hazards, the relative importance of each fish or shellfish population (and therefore hazard) differed. Each region and fleet also had its own intrinsic exposure and vulnerability profiles, further modulating the overall climate risk. However, as the base set of hazards was the same in both CRAs, a direct comparison of the two cases was possible, allowing the relative climate risk to coastal-regions and fleets to be gauged.

Systematic differences in risk between fleets and coastal communities were seen among European countries (Figure 3) and several characteristic types of responses were apparent. Countries in south-eastern Europe, together with the United Kingdom, had the highest risk across both fleets and coastal regions. The climate risk scores of regions on the south coast of the Baltic Sea (Latvia, Lithuania, Estonia and Poland) were typically higher than their fleet level scores, while the high fleet risk of NW European states was offset by their low risk to regions. Spain and Sweden were characterised by generally low climate risks in both coastal regions and fleets.

**Figure 3.**
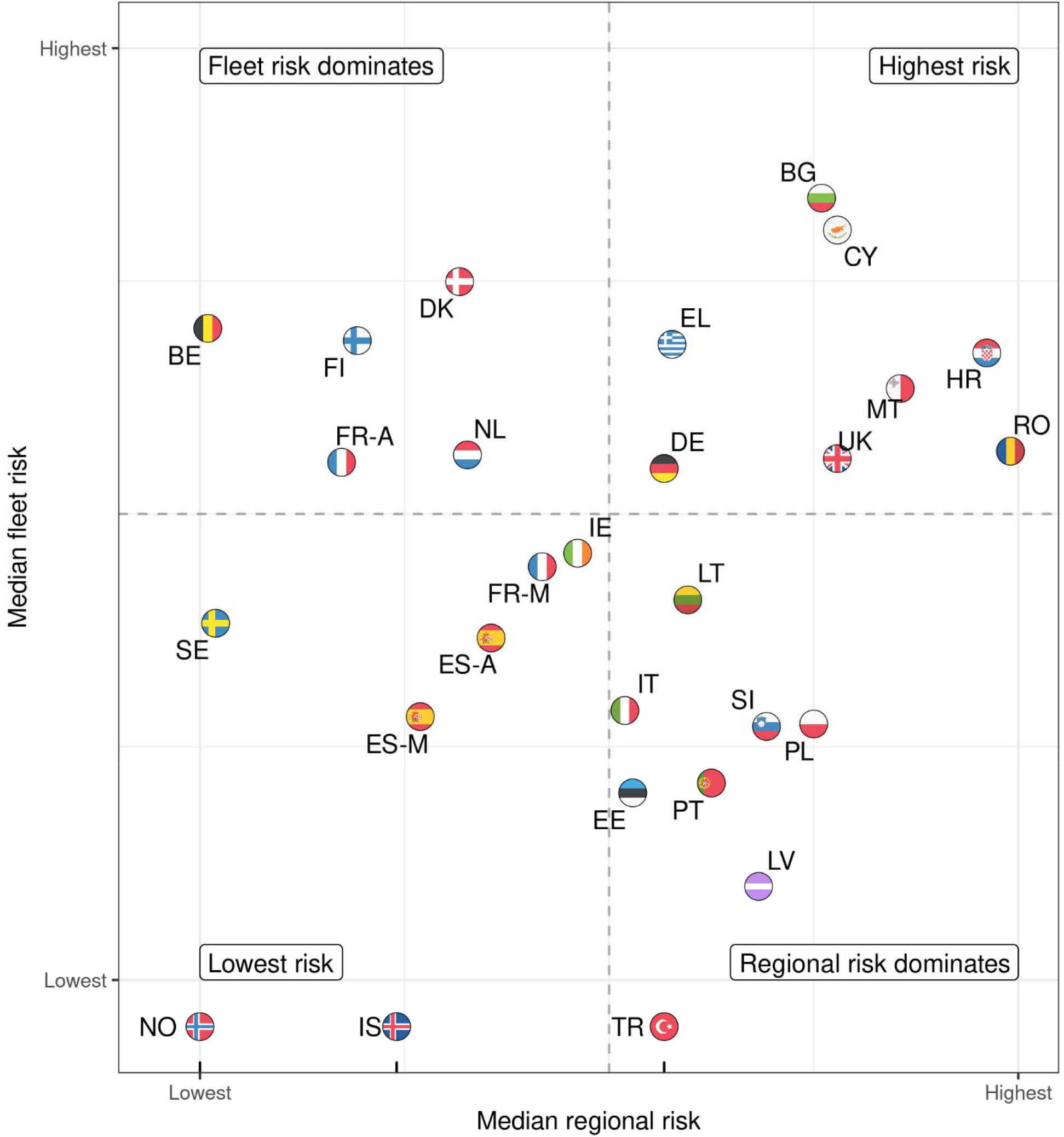
Comparison of the median fleet- and coastal-region risk rankings for European countries. Labels indicate the country code. In addition, France (FR) and Spain (ES) are split into their Atlantic (-A suffix) and Mediterranean (-M suffix) seaboards. As the fleet-segment analysis only covers fleets from the EU and UK, no data are available for Turkey, Norway and Iceland: their regional risk results are plotted in the horizontal margin. Dashed lines divide the coordinate system into quarters. Country codes: BE: Belgium. BG: Bulgaria. CY: Cyprus. DE: Germany. DK: Denmark. EE: Estonia. EL: Greece. ES: Spain. FI: Finland. FR: France. HR: Croatia. IE: Ireland. IS: Iceland. IT: Italy. LT: Lithuania. LV: Latvia. MT: Malta. NL: Netherlands. NO: Norway. PL: Poland. PT: Portugal. RO: Romania. SE: Sweden. SI: Slovenia. TR: Turkey. UK: United Kingdom.

Multiple factors act together to determine the individual risk rankings of coastal regions and fleets. In this analysis, the fishing fleets and fishery-dependent regions of any given country were both exposed to the same underlying hazard: climate change impacts on fish populations, primarily shaped by the ecology and biogeography of marine life inhabiting a country’s adjacent waters (e.g. Figure S1a). Similarly, substantial variations in biodiversity across European waters constrained the availability of species and therefore impacted the exposure score: for example, high exposure scores characterised states bordering the low-biodiverse Baltic and Black Seas (Figure S1b). Hazard and exposure were also, however, both determined by consumer preference and economic factors that shape the species targeted by fleets and landed in regions. Socio-economic factors also shaped the vulnerability dimension: the relatively high affluence of NW European states, for example, reduced their coastal risk rankings while poorer SE European states received high vulnerability scores (e.g. Figure 3, Figure S1c). Similarly, differences between northern and southern Europe in the size and number of fishing vessels could be clearly expected to influence their profitability, and therefore fleet vulnerability. The processes driving the observed risk-patterns are therefore complex and reflect the combination of patterns functioning across multiple dimensions at multiple levels.

## Discussion and Conclusions

Our analysis highlights the wide variety of challenges facing European countries in adapting their fisheries sectors to a changing climate. In some cases, such as in the southern-Baltic states, a focus on building adaptive capacity in coastal regions would be of most benefit e.g. by creating alternative employment opportunities or providing an economic ‘safety net’ through wider social measures. In other regions, fleet risks dominate and therefore increasing the efficiency, adaptive capacity and catch diversity of the fleets would appear to be a priority. Some areas, such as the UK and south-east Europe appear to require both types of intervention and therefore present the greatest adaptation challenges. It is clear that no “one-size-fits-all” solution that can be applied across all European waters or even, in some cases, across a country (e.g. the UK): climate adaptation plans, therefore, need to be tailored to these local realities.

Climate risk and vulnerability analyses can play a key role in shaping these adaptation plans. By increasing awareness of the elements that contribute most to a fleet or coastal region’s risk (Allison *et al*., 2009), CVAs and CRAs can help maximise the effectiveness of interventions given limited resources (Lindegren and Brander, 2018). Previous socio-economic linked analyses have focused on adaptive capacity (in the CVA framework) as a focal point for action (Allison *et al*., 2009; Cinner *et al*., 2013). However, the diversity of European risk profiles found here also highlights the need for adaptation actions across all components contributing to climate risk.

Ensuring sustainable management of the living marine resources upon which the sector rests is a key action for the European fisheries sector. The impacts of over-exploitation can be more important than those stemming from climate change, particularly in the heavily fished North Atlantic region (Cheung *et al*., 2018). Maintaining stocks at a higher abundance leads to increases in genetic diversity, meta-population complexity, and age structure, all of which make stocks more resilient to the challenges of a changing environment (Drinkwater *et al*., 2010; Planque *et al*., 2010). The ensuing increase in productivity and incomes also simultaneously benefits both fishing fleets and regions, generating a “win-win” effect (Bell *et al*., 2013). Fisheries scientists already have many of the tools necessary to ensure that management systems are robust to climate change and climate variability (Pinsky and Mantua, 2014), while new tools, such as seasonal-to-decadal marine ecological forecasts and early-warning systems (Payne *et al*., 2017), can potentially provide the basis for additional coping strategies (Hobday *et al*., 2018).

Diversification is a second key action to reduce climate risk. Fishing fleets and coastal regions relying on only a few species have an elevated risk of climate impacts: increasing this spread reduces (by definition) exposure and buffers fleets and regions against climate risk (Cline *et al*., 2017; Young *et al*., 2019; Pinsky *et al*., 2020). Diversification of catches and landings can take place autonomously as fishers respond to changes in the abundance and distribution of the fish they catch (Pinsky and Mantua, 2014; Lindegren and Brander, 2018). For example, changes in the distribution of fish species in waters surrounding the UK (Perry, 2005; Simpson *et al*., 2011; Baudron *et al*., 2020) have led to the development of new fisheries for squid, seabass and red mullet, amongst others (Pinnegar *et al*., 2020). CRAs such as the one presented here can also have an important role in this process by highlighting alternative species or populations with a lower climate hazard that can be targeted, thereby further reducing risk. Alternatively, reducing fisheries dependency and diversifying income sources by, for example, participating in multiple fisheries or in tourism, recreation or aquaculture has also been shown to reduce variability in income and thereby risk (Kasperski and Holland, 2013).

There are, however, barriers to diversification (Cline *et al*., 2017; Young *et al*., 2019), including knowledge, economic and governance barriers. For example, the ability to catch new species may be limited by existing quota agreements (Pinsky *et al*., 2018) such as the EU Common Fisheries Policy “relative stability” clauses, whereby the allocation of fishing quotas is fixed to reflect historical catches from 1973-1978 (Hoefnagel *et al*., 2015). Ecology can also be constraining: the limited catch diversity and therefore high exposure of fleets and coastal regions adjoining the Black and Baltic Seas, for example, arises at least in part from the naturally low biodiversity of these seas. Changing target species or fishing technologies can also be costly, creating financial barriers to diversification (Kasperski and Holland, 2013).

Governance also has a key role to play in coordinating and driving actions to reduce the vulnerability of fleets and regions. Investments and support for developing new, and switching between existing, fishing, storage, transport and processing technologies can increase the efficiency of fleet operations and therefore reduce vulnerability (McIlgorm *et al*., 2010; Bell *et al*., 2011; Pinsky and Mantua, 2014). Increasing regional development, including employment opportunities outside the fisheries sector, reduces regional vulnerability and risk (Allison *et al*., 2009; Badjeck *et al*., 2010). Furthermore, both fishing fleets and coastal regions can also potentially benefit from governance-led actions that increase the flexibility, ability to learn, social organisation and the power and freedom to respond to challenges (Cinner *et al*., 2018). Regional, national and European governments therefore have a critical role to play in helping fisheries and ocean-dependent regions to adapt to the risks presented by climate change.

Several key caveats of our analysis need to be highlighted. Our analysis focused solely on the sensitivity to ocean warming, ignoring other climate-driven processes, such as ocean acidification, deoxygenation, and changes in storminess or circulation patterns (IPCC, 2019; Pinnegar *et al*., 2019) that, while important, we view as second order effects. Spatial differences in the rates of warming across European regional seas were also not accounted for here but the range of these rates (up to 2°C by 2050) is much smaller compared to the variability in thermal safety margins across the range of some species (range up to 15°C) (Figure S4). We were unable to find European-wide data sources that quantify the relative importance of fisheries in each of our coastal regions (e.g. Natale *et al*., 2013): filling this data gap would further refine the analysis. The treatment of uncertainty in CVAs and CRAs varies greatly between studies (Hare *et al*., 2016; Spencer *et al*., 2019) but in such a semi-quantitative analysis, the choice of metrics is usually the most important aspect (Monnereau *et al*., 2017). We believe that this “structural uncertainty” (*sensu* Payne *et al*., 2016) is best addressed by focusing on a limited but transparent and readily interpretable set of indicators, rather than by quantifying uncertainties or increasing complexity. Finally, while we have considered European fisheries targeting fish stocks that span the Mediterranean Sea, we have not incorporated coastal communities in African countries that also fish on these same stocks. The relatively low GDP per capita of these communities suggests that they would have correspondingly high regional vulnerabilities and, therefore, correspondingly high climate risk profiles but it is not possible to draw robust conclusions in the absence of appropriate data sets. Nevertheless, the population-level hazards generated here (Table S7) could be readily applied to aid such analyses in the future.

The limitations of the CRA approach itself also need to be borne in mind. Here, we explicitly focus on the “negative” impacts of a warming climate, following the IPCC framing of climate hazard in terms of “adverse effects” (Oppenheimer *et al*., 2014; IPCC, 2019), even though there will also undoubtedly be some “positive” effects (eg Kjesbu *et al*., 2014) that may partially offset the negative effects at some localities. However, estimating a “net” hazard requires quantifying changes and placing them on a common basis (e.g. economic impacts), a challenging task best done in explicitly quantitative settings such as end-to-end models rather than CRAs. Similarly, CRAs are not intended to make quantitative “predictions” of the future (Pecl *et al*., 2014) and we are not aware of cases where they have been used in this manner. Rather, the strength of the CRA approach is its ability to provide a transparent and consistent framework for the rapid assessment of climate risk across a (very) wide range of fleets and coastal regions, allowing the most at-risk elements to be identified and prioritised for adaptation actions (Pinnegar *et al*., 2019). Developing climate adaptation plans for local situations should involve detailed quantitative modelling to provide the necessary details.

This study has shown that, even though the average climate risk to European countries is moderate compared to many other countries across the globe (Allison *et al*., 2009; Blasiak *et al*., 2017), major differences exist across the European continent. This corroborates with fine-scale spatial differences among fishing communities documented in eastern North America (Colburn *et al*., 2016; Rogers *et al*., 2019) and the Caribbean (Monnereau *et al*., 2015; Pinnegar *et al*., 2019), where individual communities would be best served by different adaptation actions. Our detailed analyses allow a distinction between climate hazard, exposure and vulnerability as key sources of climate risk to fleets and coastal regions and highlight where (and what) adaptation measures can have the greatest impact in increasing resilience, given limited financial resources.

## Acknowledgements

This project received funding from the European Union’s Horizon 2020 research and innovation programme under grant agreement No 678193 (CERES – Climate change and European Aquatic Resources). The results generated by this analysis can be explored using an online tool available at https://markpayne.shinyapps.io/CERES_climate_risk/ Source code is available at https://github.com/markpayneatwork/CERES_vulnerability. “Fishing Boat”, “Urban” and “Thermometer” icons in Figure 4 by smalllikeart from www.flaticon.com.

**Figure 4.**
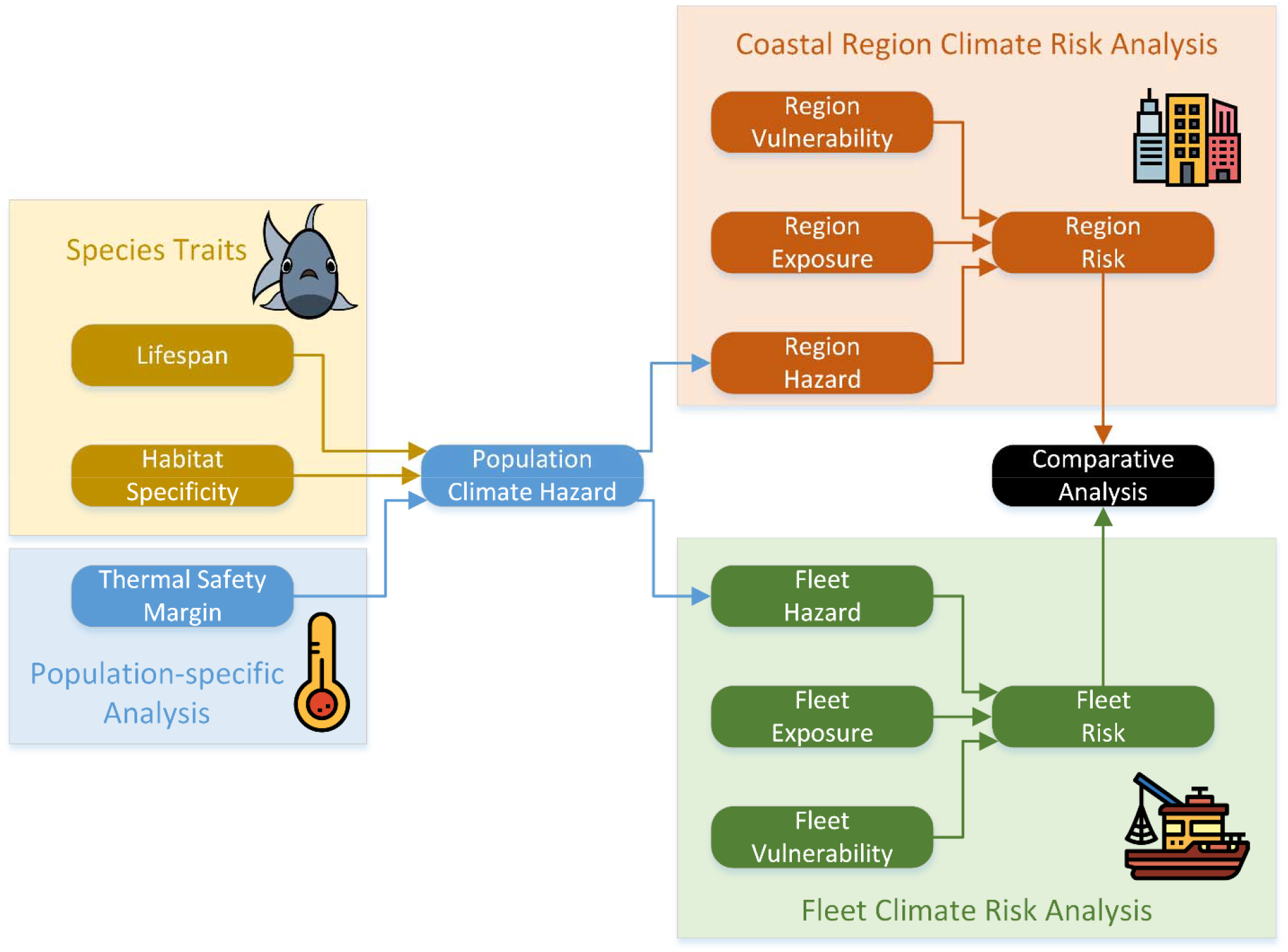
Schematic diagram illustrating the approach used here to estimate climate risk in European fishery-dependent coastal regions and fishing fleets. Species traits and population specific analyses of the thermal safety margin are combined to give a population-specific climate hazard. This hazard then forms the basis for the region and fleet level CRAs, based on the combination of hazard, exposure and vulnerability. Finally, the region and fleet risks are combined again into a comparative analysis. A detailed flow diagram is presented in the supplementary material (Figure S3).

## Methods

### General approach

We have applied an integrated approach to a climate risk assessment (CRA) across the European fisheries sector. The CRA has three major components (Figure 4; Figure S3). The first and most fundamental of these is the *population hazard* component, where the hazard associated with adverse climate change impacts on individual fish populations is quantified. We then use these hazard metrics as inputs into two parallel climate risk assessments focussing on coastal regions and fishing fleets in turn. In each of these cases, the population hazard is integrated up to the region or fleet level based on information about the relative importance of each fish population to that unit to form the region- or fleet-specific hazards. These hazard data are then complemented with region- and fleet-focused *exposure* and *vulnerability* metrics to produce a *climate risk* for each. Finally, we combine the risks from each component into a comparative analysis across nations.

### Scope and Data Sources

We aimed to assess the climate risk for the European marine fisheries sector, including all 22 EU countries with marine borders, the United Kingdom, Norway, Iceland and Turkey. We based our analysis primarily on catch data from FAO Areas 21, 27, 34 and 37 held in the EUROSTAT database (Table S1), excluding distant water fleets. While this database covers more than 1200 species, many of these are economically minor. We therefore simplified our task by focussing on species making up the largest 90% of the value of the marine fish and shellfish sector in each country and across Europe as a whole. Two species predominately inhabiting freshwater, European perch (*Perca fluviatilis*) and pike-perch (*Sander lucioperca*), were removed from the database. Misspelled (or alternative) scientific names were corrected where we could identify these (following World Register of Marine Species, WoRMS) (Table S3).

Regional analyses were performed for European coastal regions based on NUTS2 statistical units. Sub-national indicators of landings composition were derived from monthly harbour-level “first-sales” data from the EU Market Observatory for Fisheries and Aquaculture (EUMOFA) (Table S1). In cases where this data covered more than one NUTS2 unit within a country (10 countries), the harbour data was aggregated up to NUTS2 units based on the geographical coordinates of the harbours. Where EUMOFA data coverage was insufficient, the coastal NUTS2 units of that country were merged into one “region” (Table S5) and EUROSTAT national landings data were assigned to it (Table S1). Socio-economic data for the NUTS2 units was also obtained from EUROSTAT and integrated up to our “regions”, if relevant.

The Annual Economic Report (AER) provided by the EU Scientific, Technical and Economic Committee for Fisheries (STECF) (STECF, 2018) formed the basis of the fishing fleet analysis (Table S1). This dataset has the advantage of providing a single coherent source for fleet segments (the combination of fishing technique and a vessel length category) across all of the European Union and United Kingdom: however, it does not include data on fleets from Norway, Iceland or Turkey, and in the absence of comparable datasets, these countries were not included in this part of the analysis.

All data were averaged over the period 2010-2018, where available.

### Hazard Metrics

The *hazard* dimension of our CRA measures the strength and severity of adverse climate change impacts on the unit of interest: in this case, fish populations in European waters. Many previous CVAs and CRAs do not distinguish between the positive and negative effects of climate change, and simply highlight elements of their study system that will change, making interpretation difficult. In contrast, and following the IPCC’s definition of risk in relation to an “adverse event” (IPCC, 2019), we focus explicitly on “negative” impacts in order to have an unambiguous interpretation.

We consider the hazard due to climate change impacts on living marine resources as being the combination of both species-specific and population-specific processes as follows:

### Species-specific processes

A trait-based approach was employed to characterise the hazard of climate change to a species. Such an approach is well established in climate risk and vulnerability analyses (Pecl *et al*., 2014; Hare *et al*., 2016; Jones and Cheung, 2018), due to its ability to draw on general understanding of the response of species to climate change. Trait data was collated from previously published databases (Engelhard *et al*., 2011; Pecuchet *et al*., 2017; Beukhof *et al*., 2019b, 2019a) and complemented with data from Fishbase (Froese and Pauly, 2019) and Sealifebase (Palomares and Pauly, 2019) (accessed April-July 2019) (Table S1). Of the original set of species from EUROSTAT, 24 taxa were only at the genus level, and appropriate trait sets were therefore identified based on ‘exemplar species’: in some cases different exemplar species were used for the North Atlantic (FAO Area 27) and Mediterranean regions (FAO Area 37) (Table S2). Barnacles (*Pollicipes pollicipes*) and solen razor clams (*Solen* spp.) were removed from the analysis owing to a lack of comparable biological trait data and difficulties identifying suitable exemplar species.

Trait selection aimed to avoid double-counting information due to inclusion of correlated traits, a commonly overlooked issue (Pecuchet *et al*., 2017) that impacts many published analyses (Cheung *et al*., 2005, 2018; Hare *et al*., 2016; Jones and Cheung, 2018). For example, smaller fish are typically planktivorous, live shorter and grow faster, giving a high correlation between maximum length, lifespan, growth rates and trophic level. Lifespan is the most commonly available of these metrics and was therefore chosen as an exemplar for this set of traits. Shorter lifespans are associated with seasonal and variable environments (Pecuchet *et al*., 2017), implying robustness to change and variability, paralleling the approach used in other studies (Cheung *et al*., 2005, 2018; Hare *et al*., 2016; Jones and Cheung, 2018).

A “habitat specificity” metric was also developed. Species with spatially restricted habitat requirements during part or all of their life-history are recognised as being more sensitive to disruption (Rijnsdorp *et al*., 2009; Petitgas *et al*., 2013). In addition, mobile species have the ability to move rapidly to avoid unfavourable conditions in a way that sedentary species do not, and therefore have a lower climate hazard (Pinnegar *et al*., 2019). Traits defining the mobility and vertical and horizontal habitats were therefore collated into a single “habitat-specificity score” (Table 2). The final set of traits is included as supplementary material (Table S6).

**Table 2.**
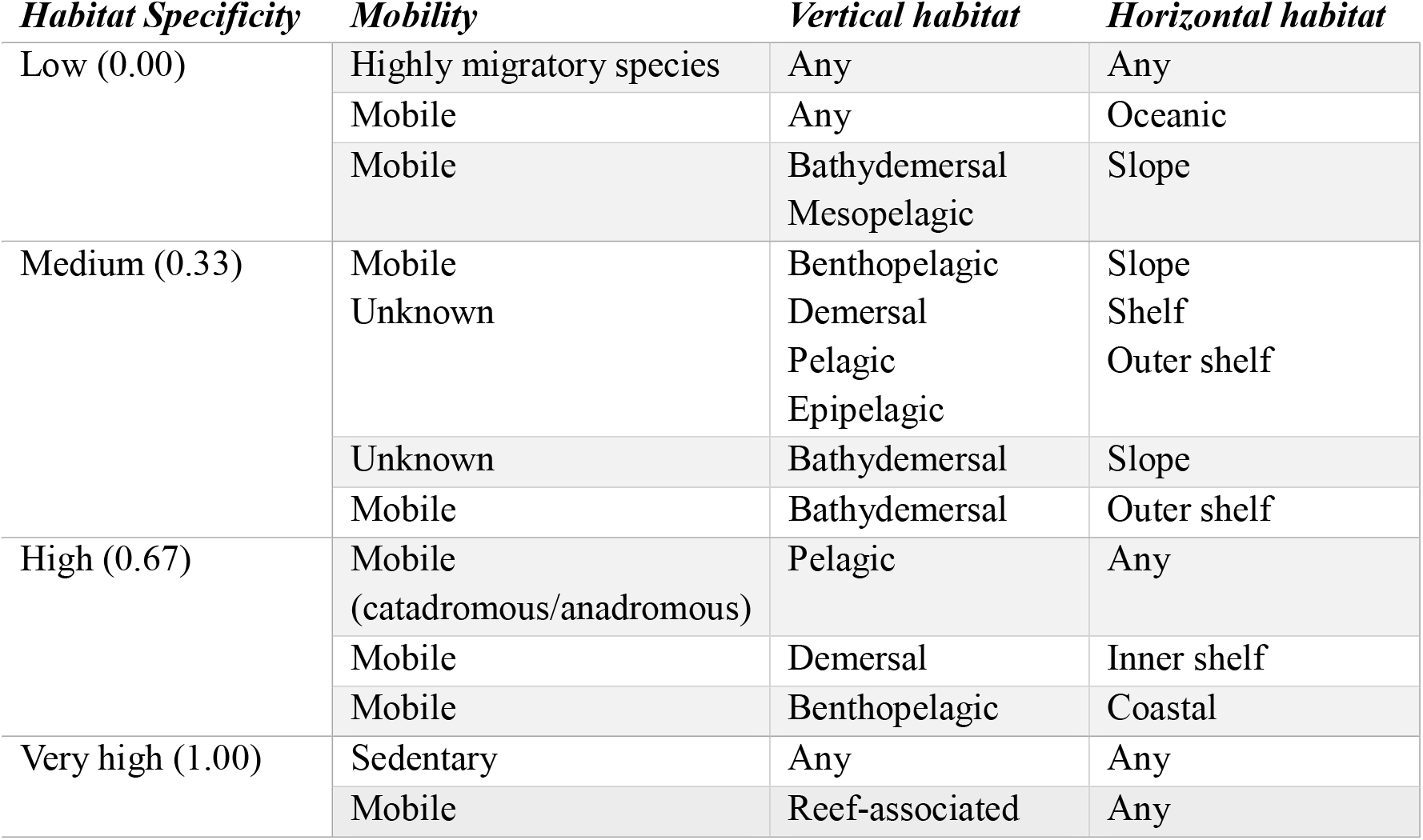
Combination of mobility, vertical and horizontal habitat traits to generate a habitat specificity score. Trait categories follow the scheme of Engelhard *et al* (Engelhard *et al*., 2011).

### Population-specific processes

The stress a fish population experiences as the ocean warms depends on the amount of warming, a commonly employed metric of exposure in CVAs (Allison *et al*., 2009; Hare *et al*., 2016). However, the physiological context of this warming is also critical but often overlooked. For example, cod (*Gadus morhua*) in the North Sea are close to their upper thermal limit, and will therefore experience negative impacts of warming, while cod in the Barents Sea are far from this limit and will experience little or no negative effects of the same amount of warming (Drinkwater, 2005). Such a spatial and physiological context of warming, often overlooked in many CRAs and CVAs, is critical to differentiate the climate hazard among different populations of the same species.

We resolve this problem in two ways. First, we perform our analysis at the “population” level, defined as the combination of species and FAO subarea e.g., cod in subarea 27.4 (North Sea). Note that while this approach is similar to that used to manage many European fish stocks, we explicitly avoid the use of the term “stock” to refer to this unit of analysis, as it has clear implications in fisheries management but is not always the same as our definition “population”. Populations comprising less than 5% of the total catch of the species were excluded from the analysis. Secondly, we place the degree of warming experienced by these populations in a physiological context using estimates of the thermal-safety margin (TSM) (Deutsch *et al*., 2008; Sunday *et al*., 2014; Pinsky *et al*., 2019; Dahlke *et al*., 2020). The TSM is estimated as the difference between the empirically-derived maximum temperature that the species prefers and the temperature of the environment: high TSMs indicate a high capacity to tolerate warming. Population-specific TSMs therefore permit a fine-grained measure of the warming-related hazard.

We derived population-specific estimates of TSM from the habitat models, parameters and maps provided by Aquamaps www.aquamaps.org (Kesner-Reyes *et al*., 2012) (Table S1). We downloaded “native distribution maps” from the Aquamaps website for the species included in our analysis: where multiple maps were available, choice was guided by Aquamaps’ internal quality ranking system. For the invasive species veined rapa whelk (*Rapana venosa*), originally from waters around Japan, Korea and China but now supporting a large fishery in the Black Sea, the “Suitable Habitat map” was used. From each species’ map we used the “90^th^ percentile” parameter for the temperature response as an empirical estimate of its upper thermal tolerance. Temperatures in a subarea were based on the data underpinning the Aquamaps model (NOAA NCEP Climatology, 1982-1999) (Kesner-Reyes *et al*., 2012), ensuring congruence between the tolerance parameters and the temperature data. Sea-surface or -bottom temperature data, as used in generating the species’ Aquamap, were masked using the habitat model to eliminate unsuitable habitat for each individual species (Figure 5). Projected temperatures changes from 1999 to 2050 under the SRES A2 scenario were also available in this dataset and extracted for each population in the same manner for use in supporting analyses (Figure S4). Population-specific estimates of TSM were calculated as the median difference between the species’ “90^th^ percentile” parameter and temperature across all valid pixels in that subarea.

**Figure 5.**
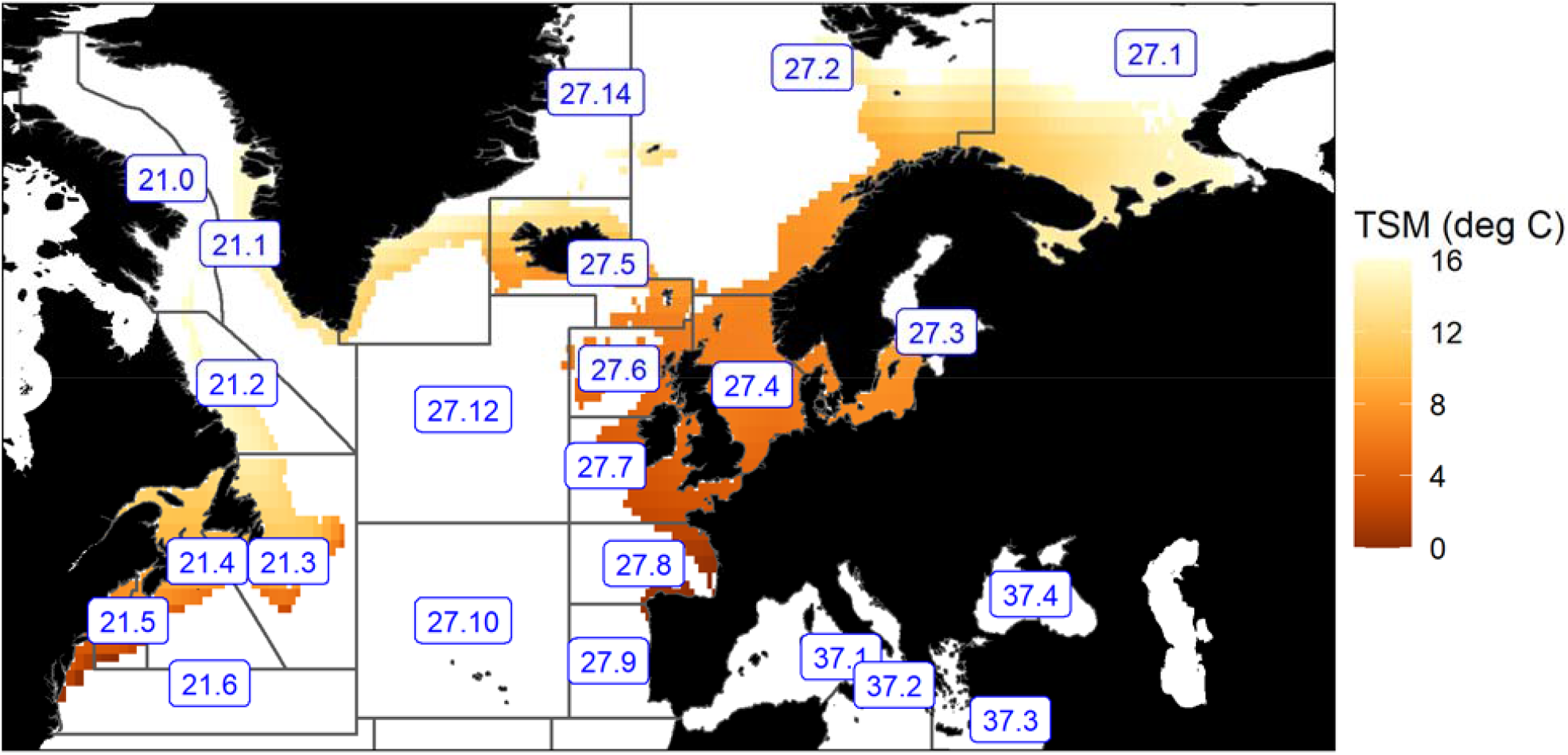
Use of Aquamaps to calculate TSM metrics. Atlantic cod (*Gadus morhua*) as an example. Environmental data and species thermal tolerance data from Aquamaps are used to estimate the thermal safety margin (TSM) for this species (coloured pixels) and masked using the habitat model to limit data to modelled regions of occurrence. Median TSM values are then calculated within each FAO subarea defining a population (grey polygons, blue labels).

### Population-level hazard

Hazard metrics were combined based on their relative ranking for each population. We chose to give equal weight to the species (lifespan, habitat-specificity) and population-level (TSM) aspects of the analysis when combining the metrics: after converting to a rank percentile, a weight of 0.25 was given to the species’ lifespan (shorter-lifespans give a low hazard), 0.25 for the species’ habitat-specificity (low specificity gives a low hazard) and 0.5 to the population TSM (high TSMs give a low hazard). Equal weighting of the metrics (0.33 / 0.33 / 0.33) was also considered (Hamilton and Ferry, 2018) but the resulting hazard metrics were found to be strongly correlated with the original (0.25 / 0.25 / 0.50) weighting (Spearman correlation coefficient of 0.95; Figure S5), indicating that the relative hazard ranking of individual populations under the two schemes is very similar.

Population-level hazard scores were integrated up to coastal region and fishing-fleet levels. In the case of the fleet analysis, this was based on the relative composition (by value) of the populations that each fleet fishes on, while in the case of the coastal region analysis it was based on the composition (by value) of landings in that region (Figure 4, Figure S3).

### Exposure metrics

We define exposure as an indicator of how sensitive a coastal region or fishing fleet is to changes in the fish populations it is dependent on. Fleets or coastal regions have lower exposure if they catch a wide range of different fish species, rather than concentrating on a specific resource (Cline *et al*., 2017; Pinnegar *et al*., 2019; Young *et al*., 2019). If one species is reduced or lost due to the effects of climate change, the impact of that loss is relatively less severe for fleets and coastal regions that are dependent on a broad portfolio of species. We therefore defined our exposure metrics following this logic, using two different metrics to characterise diversity of catch or landings: i) the Shannon diversity index, one of the most commonly used diversity indices in ecology and ii) Simpson’s dominance index, a statistic that emphasizes the relative abundance of the most common species in the sample (Pinnegar *et al*., 2019).

For coastal regions, exposure metrics were based on the value of landings data from EUMOFA and EUROSTAT (Table S1; Figure S3). While EUROSTAT data is species resolved, EUMOFA data is organised in approximately 100 “main commercial species” (MCS) groupings: we therefore harmonised the two datasets by aggregating EUROSTAT data to the MCS groupings based on correlation keys provided by EUMOFA. The Shannon and Simpson metrics were then calculated to estimate the diversity of MCS groups.

For fleet segments, the value of landings is available by species code from the STECF Annual Economic Report (STECF, 2018). The two diversity indices could therefore be calculated directly to quantify the diversity of species caught.

In both cases, the exposure index was produced as a composite index of the two indices described above by averaging the percentile ranks and then re-calculating percentile ranks again.

### Vulnerability metrics

*Vulnerability* in this setting refers to the ability of the analysis unit (either a coastal region or a fleet) to effectively address the hazard via adaptation or coping strategies.

The regional vulnerability metric was based on the gross-domestic product per capita of the region, as calculated from EUROSTAT data at the NUTS2 level (Table S1). Regions with high GDP per capita were viewed as having a high adaptive capacity and therefore low vulnerability. Regional vulnerability was calculated as the percentile rank of this statistic.

Fleet segment vulnerability was based on the net profit margin (NPM). This is a standard economic metric, defined as net profit (i.e. revenue minus variable, fixed and opportunity costs) divided by the total revenue: it therefore represents how much of the total income generated by the fleet is net profit (STECF, 2018). NPM takes into account many of the different factors that influence the profitability of the fleet, and is also scale independent (as profitability is divided by the revenue), allowing comparison of both large and small segments. NPM was calculated for each fleet segment based on economic data from the STECF Annual Economic Report (STECF, 2018) (Table S1), and the vulnerability score generated based on percentile rank. Fleet segments with high profitability were viewed as being less vulnerable to the effects of climate change, as they could absorb the potential loss associated with a climate change having a negative impact on their target species.

### Climate risk metrics

For each of the coastal regions, and for each of the fleet segments, the overall *climate risk* was calculated as the unweighted mean of the hazard, exposure and vulnerability percentile ranks.

