## Supplementary Figures for "Climate risk to European fisheries and coastal communities"

Corresponding author: Mark R Payne

**This PDF file includes:**

Figures S1 to S5

**Other supplementary materials for this manuscript include the following:**

Datasets S1 to S9

**Figure S1 Components of climate risk by coastal region**. a) Climate hazard, b) exposure and c) vulnerability are plotted for each coastal region. Colour scale is linear in the ranking but is presented without values, as they have little direct meaning. National-level borders are shown for reference. Inset at bottom-left shows small regions.

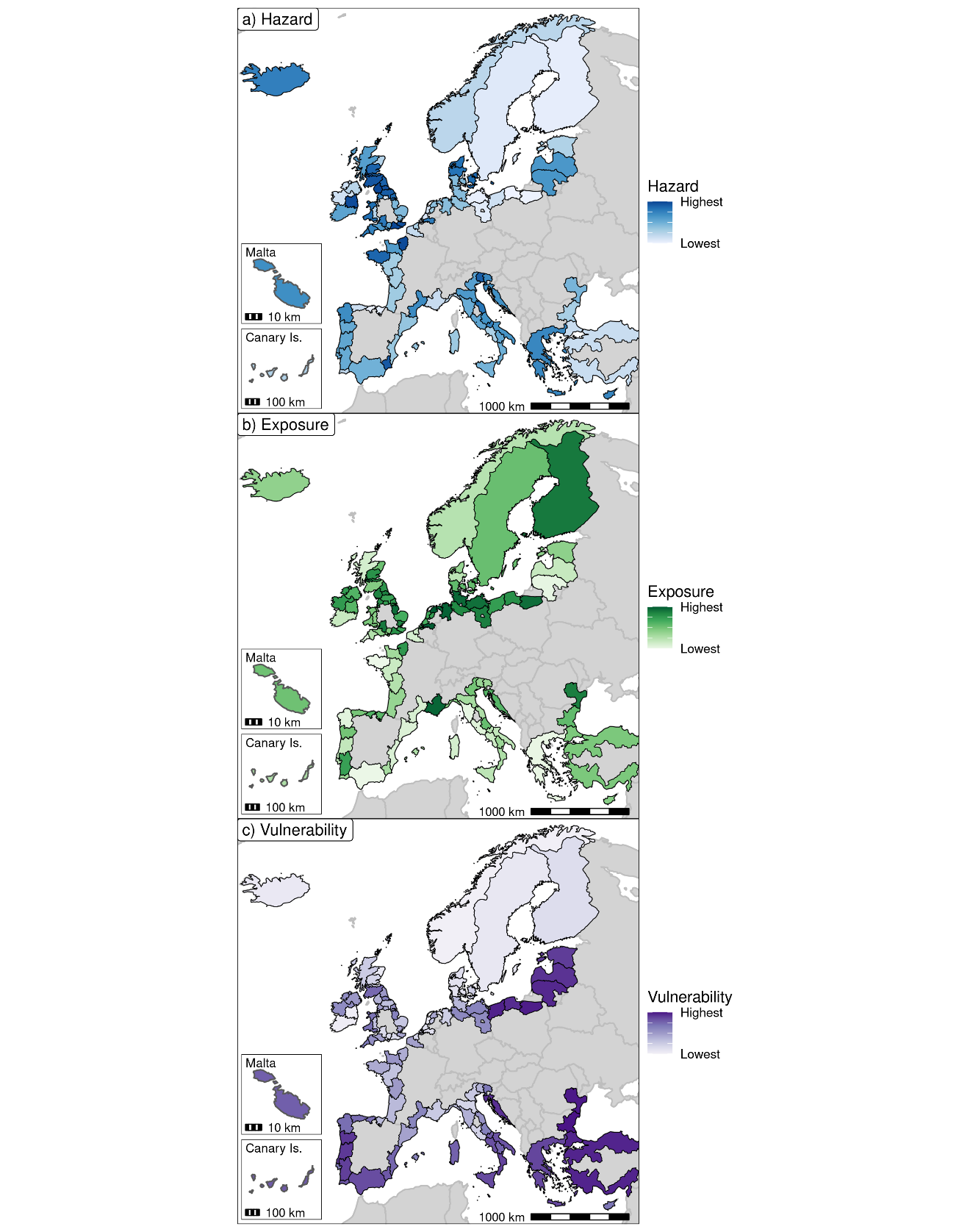

**Figure S2 Median climate risk to fishing fleets in the EU and UK**. Colour scale is linear in the ranking of the risk but is presented without values, as they have little direct meaning. National-level borders are shown for reference. Inset at bottom-left shows small regions.

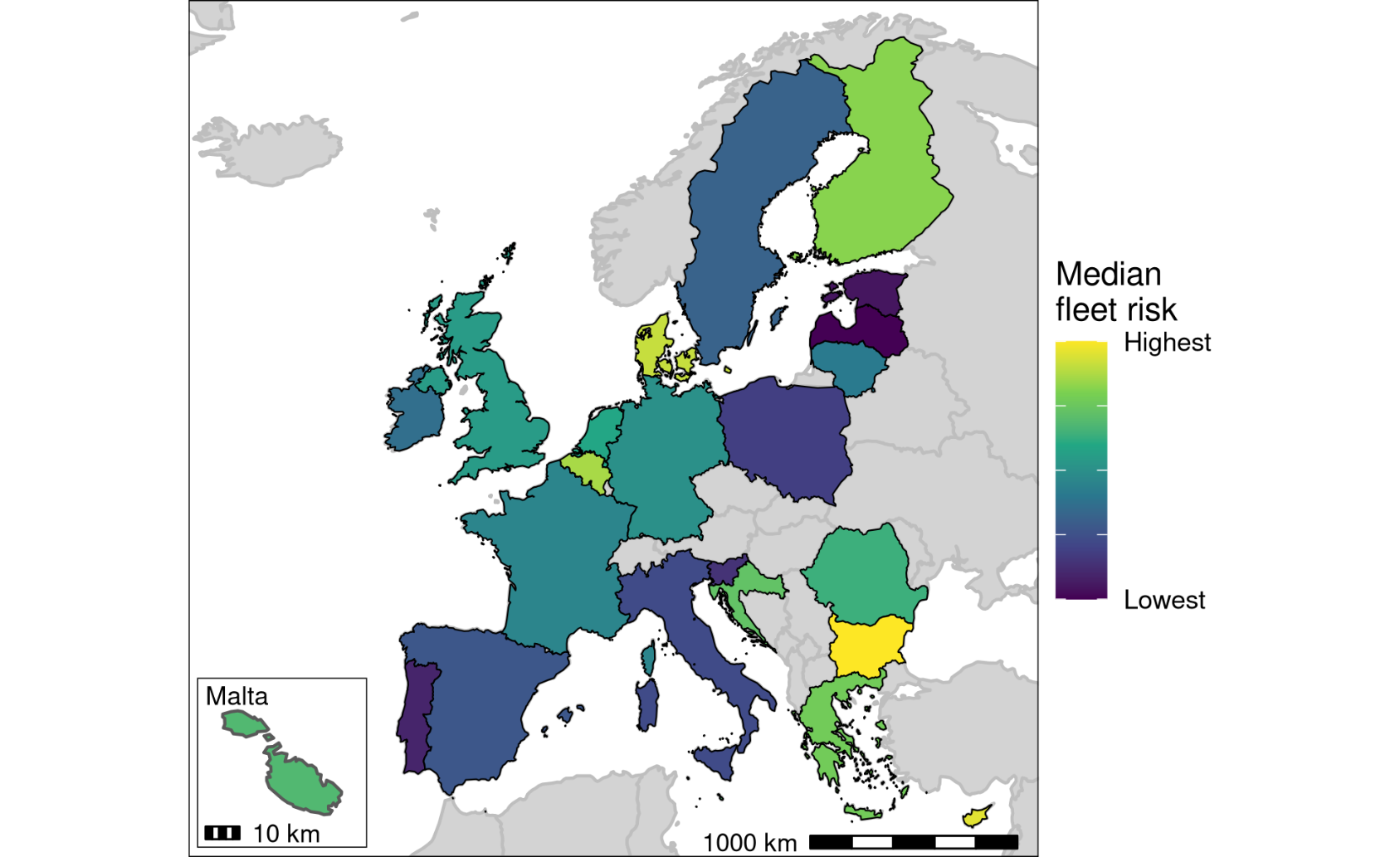

**Figure S3 Detailed flow diagram showing the relationship between datasets in the analysis.** Miniatures of the figures presented in the manuscript are included at the appropriate points in the flow diagram. The general flow of information is from left to right. External data sources are marked with heavy black borders.

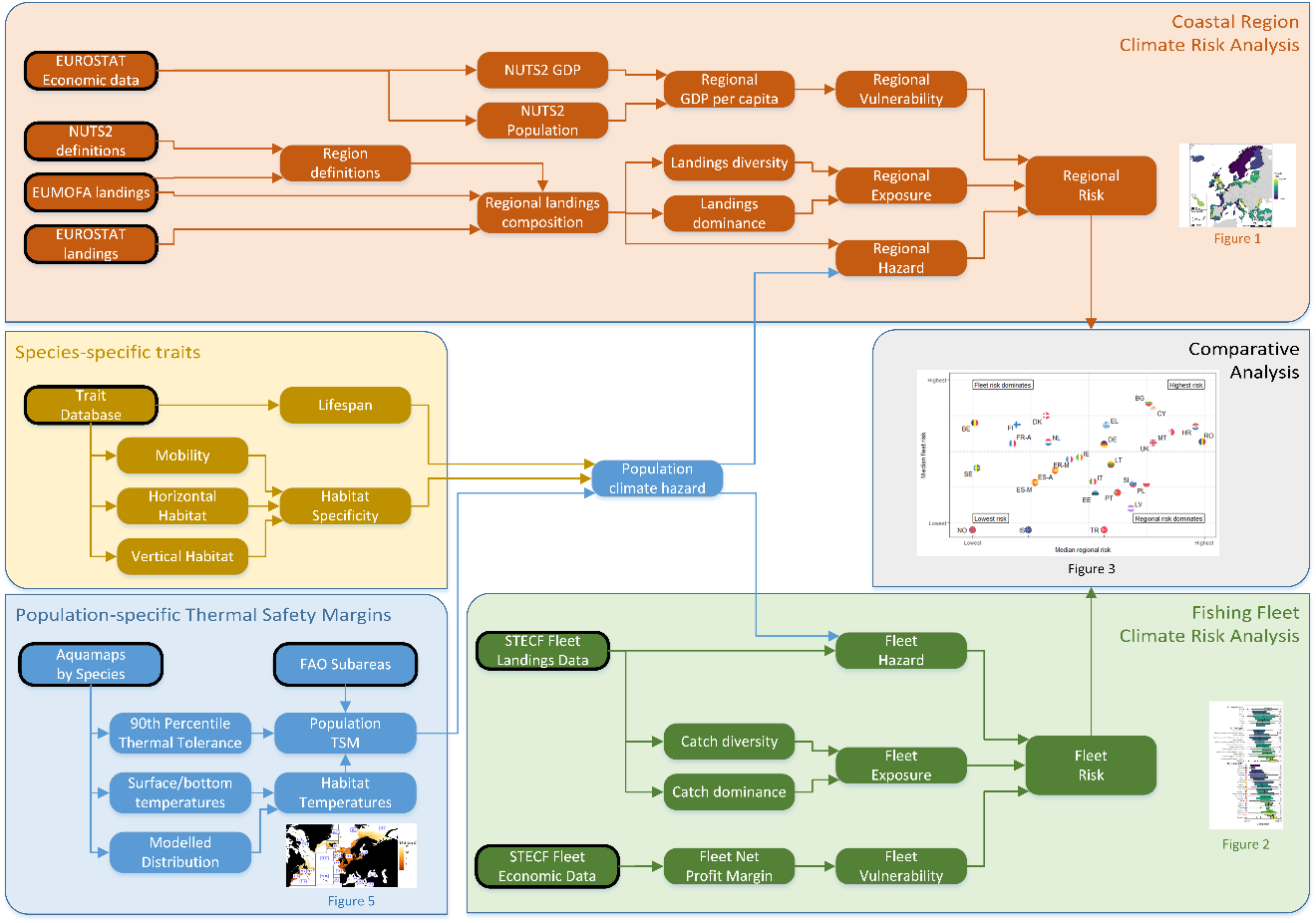

**Figure S4 Comparison of distributions.** The distribution of the projected warming over the period 1999-2050 (upper curve) and the thermal safety margin, TSM (lower curve) is shown across the populations considered in the analysis. Both curves are scaled to have the same area.

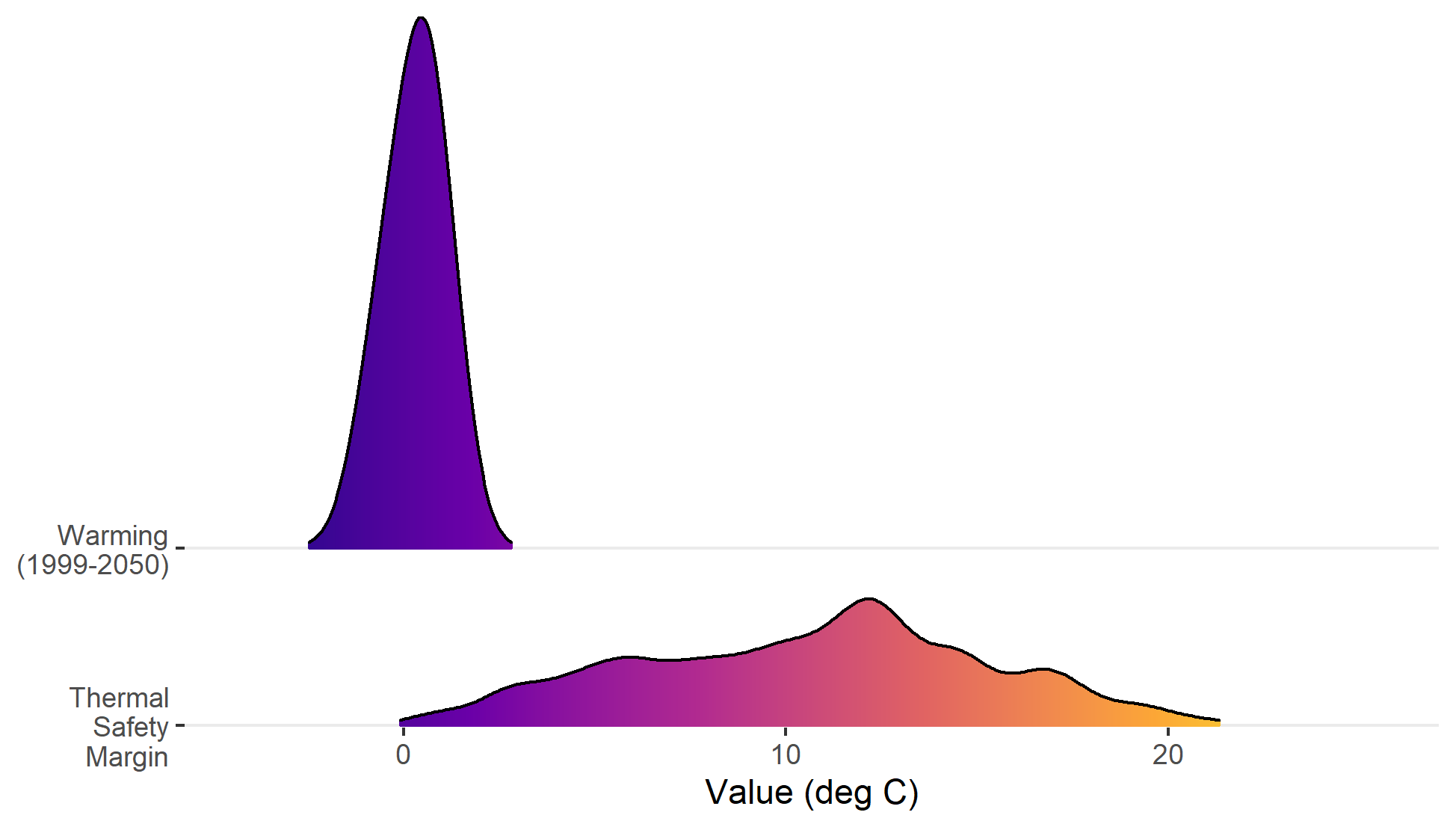

**Figure S5 Effect of alternative trait-weighting schemes on the climate hazard.** The Spearman correlation coefficient between the climate hazard using alternative weightings of the habitat specificity, lifespan and thermal safety margin (TSM) (corresponding to the three axes) and that using the current weighting is show as binned colours. The current weighting scheme (TSM = 0.50, Habitat =0.25, Lifespan = 0.25) is shown (filled circle), while a selected alternative weighting scheme giving equal weight to each component (0.33 / 0.33 / 0.33) is also shown (filled triangle). The constraint that weightings must add to 1 is reflected in the use of a ternary plot (68): dashed lines indicate the coordinates of the two highlighted points.

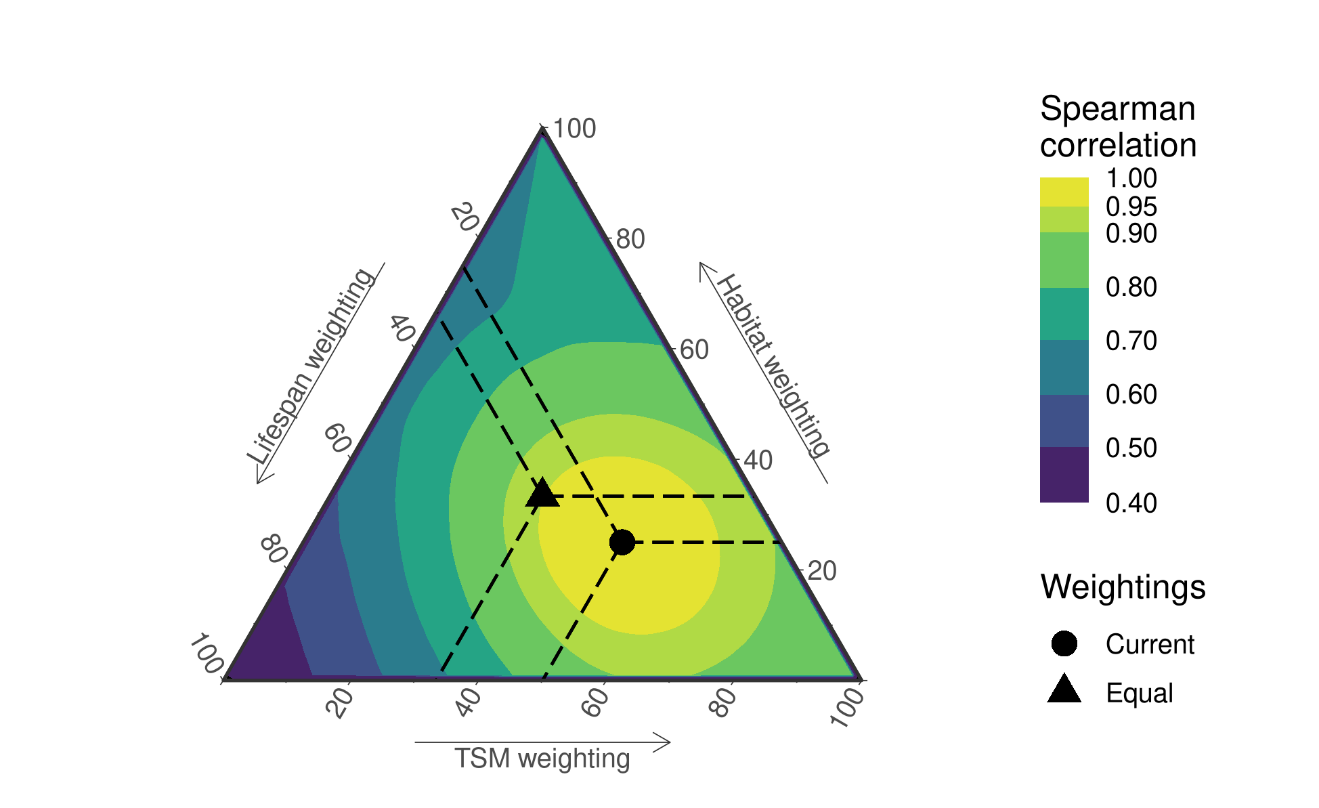
